# Features of ChIP-seq data peak calling algorithms with good operating characteristics

**DOI:** 10.1101/037473

**Authors:** Reuben Thomas, Sean Thomas, Alisha K Holloway, Katherine S Pollard

**Affiliations:** Gladstone Institutes, San Francisco, CA 94158, USA; Division of Biostatistics, University of California, San Francisco, CA 94158, USA; Phylos Biosciences, Portland, OR 97201, USA; Institute for Human Genetics and Institute for Computational Health Sciences, University of California, San Francisco, CA 94158, USA

## Abstract

**Author description:** Reuben Thomas is a Staff Research Scientist in the Bioinformatics Core at Gladstone Institutes

Sean Thomas is a Staff Research Scientist in the Bioinformatics Core at Gladstone Institutes

Alisha K Holloway is the Director of Bioinformatics at Phylos Biosciences, visiting scientist at Gladstone Institutes and Adjunct Assistant Professor in Biostatistics at the University of California, San Francisco.

Katherine S Pollard is a Senior Investigator at Gladstone Institutes and Professor of Biostatistics at University of California, San Francisco.

**Key Points:**

- Peak-calling using Chip-seq data consists of two sub-problems: identifying candidate peaks and testing candidate peaks for statistical significance.
- Twelve features of the two sub-problems of peak-calling methods are identified.
- Methods that explicitly combine the signals from ChIP and input samples are less powerful than methods that do not.
- Methods that use windows of different sizes to scan the genome for potential peaks are more powerful than ones that do not.
- Methods that use a Poisson test to rank their candidate peaks are more powerful than those that use a Binomial test.

**Abstract:** Chromatin immunoprecipitation followed by sequencing (ChIP-seq) is an important tool for studying gene regulatory proteins, such as transcription factors and histones. Peak calling is one of the first steps in analysis of these data. Peak-calling consists of two sub-problems: identifying candidate peaks and testing candidate peaks for statistical significance. We surveyed 30 methods and identified 12 features of the two sub-problems that distinguish methods from each other. We picked six methods (GEM, MACS2, MUSIC, BCP, TM and ZINBA) that span this feature space and used a combination of 300 simulated ChIP-seq data sets, 3 real data sets and mathematical analyses to identify features of methods that allow some to perform better than others. We prove that methods that explicitly combine the signals from ChIP and input samples are less powerful than methods that do not. Methods that use windows of different sizes are more powerful than ones that do not. For statistical testing of candidate peaks, methods that use a Poisson test to rank their candidate peaks are more powerful than those that use a Binomial test. BCP and MACS2 have the best operating characteristics on simulated transcription factor binding data. GEM has the highest fraction of the top 500 peaks containing the binding motif of the immunoprecipitated factor, with 50% of its peaks within 10 base pairs (bp) of a motif. BCP and MUSIC perform best on histone data. These findings provide guidance and rationale for selecting the best peak caller for a given application.

## Introduction

Regulation of gene expression is one of the fundamental means by which cells adapt to internal and external environments. Many regulatory mechanisms rely on modifying or ’marking’ the DNA in particular ways, either through covalent modification or by intermolecular interactions. ChIP-seq data are generated to provide readouts of these modifications, such as the location and intensity of binding of a transcription factor or the distribution of histone modifications that are used by the cell to establish or maintain specialized chromatin domains.

The data for ChIP-seq peak calling are stacks of aligned reads across a genome. Some of these stacks correspond to the signal of interest (e.g. binding of a transcription factor or modified histone). Many other stacks are regarded as molecular or experimental noise, or as being influenced by a systematically greater accessibility of measurement by the experiment at that particular genomic location. This manuscript deals with the problem of separating signal from noise in the stacks of reads to estimate where the immunoprecipitated protein is bound to DNA.

Many methods target this problem. The newer methods make claims of superiority over a subset of the existing ones by displaying performance on certain metrics. However, the generalizability of these performance results is unclear given the number of data sets (typically 3-5), methods, parameter settings (often only default settings), and performance metrics employed. There have been efforts to benchmark peak calling methods [1-5]. These have the advantage of being independent evaluations, but they have also been limited in scope and some times provide conflicting advice due to differences in numbers and widths of peaks evaluated [6]. For example, Harmanci et al [7], Koohy et al [5], and Micsinai et al [4] disagree about the ranking of SICER [8], F-seq [9] and MACS [10]. Consequently, the field lacks systematic recommendations for calling peaks in different scenarios.

We address these criticisms of benchmarking efforts by first abstracting the peak-calling problem into two sub-problems, identifying peaks and testing peaks for significance, respectively. We identify four features of the algorithms that differentiate them in terms of how they address the first sub-problem and eight features that differentiate the algorithms on the second sub-problem. We pick six methods that span most of the possible feature values for the two sub-problems. We then simulate ChIP-seq data for transcription factor binding to generate 100 independent ChIP and input sample pairs at each of three different noise levels to evaluate the operating characteristics of the six methods by varying the respective thresholds for determining peak significance. We also test the methods on data from one transcription factor and two histone mark experiments. Combining results across these data with mathematical analyses, we determine which features of peak-calling methods optimize performance.

## Methods

### Choice of peak calling methods

We surveyed 30 methods[4,7,9-32] in the literature and annotated each of them in terms of how they solve sub-problems 1 and 2 (see Tables 1 and S1). We chose MACS2[10], MUSIC[7], GEM[13], ZINBA[11], BCP[14] and a Threshold-based method (TM) for further analysis to balance the need to cover most of the feature space of the different methods with that of directly testing a feasible number of the more recent and/or popular methods (Supplementary Text).

**Table 1.** Features of peak-calling methods. See Table S1 for a more complete list of features and peak-calling methods.

|  | GEM | BCP(TF) | BCP(Histone) | MUSIC | MACS2 | ZINBA | TM |
| --- | --- | --- | --- | --- | --- | --- | --- |
| **Locating the potential peaks** |  |  |  |  |  |  |  |
| High Resolution | Yes | Yes | No | Yes | Yes | No | Yes |
| ChIP and input sample signals combined | No | No | No | No | No | Yes | Yes |
| Multiple alternate window sizes | Yes | Yes | Yes | Yes | No | No | No |
| Use of variability of local signal | Yes | Yes | Yes | No | Yes | Yes | No |
| **Ranking of peaks** |  |  |  |  |  |  |  |
| Binomial test | Yes | No | No | Yes | No | No | No |
| Poisson test | No | Yes | No | No | Yes | No | No |
| Normalized difference score | No | No | No | No | No | No | Yes |
| Use of underlying genome sequence | Yes | No | No | No | No | No | No |
| Posterior probability of enrichment | No | No | Yes | No | No | Yes | No |

### Simulations

ChIP-seq data representing transcription factor binding and corresponding input samples are simulated using functions adapted from the ChIPsim [33] bioconductor [34] package in R[35] that are in turn based on [36]. The data are simulated for pairs of ChIP and corresponding input samples under three noise settings - high noise, medium noise and low noise (details in Supplementary Text).

To ensure that our simulated ChIP and input data resemble real ChIP-seq experiments, we compared them to data from the first 10 million base pairs (bp) of chromosome 1 in a ChIP-seq experiment on the transcription factor Tbx5 in mouse cardiomyocytes (Luna-Zurita et al. Cell 2016). Peaks were identified using MACS2, which is one of the best performing methods on the simulated data. Input was quantified using reads per 1000-bp window. Quantiles of simulated ChIP and input data match the real data very well (Figure 1), and browser tracks resemble real data visually (Figure S1). The only difference detected is a more extreme number of reads in about 1% of simulated regions, both ChIP and input, compared to the real data.

**Figure 1.**
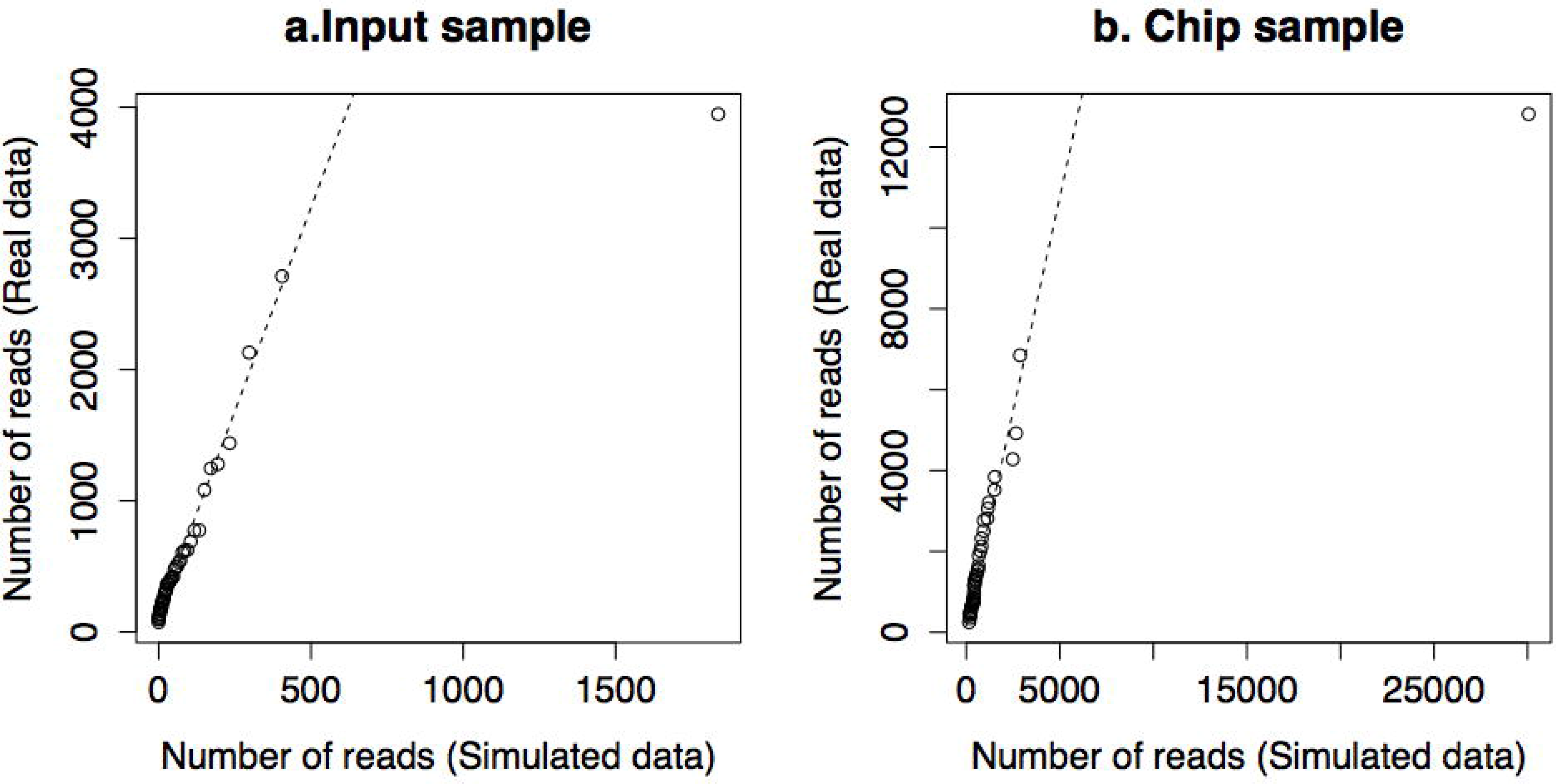
Quantile-quantile plots comparing the distributions of reads in the input and ChIP sample in one of the simulated data sets with those in a real Tbx5 ChIP sample from mouse cardiomyocytes

### Evaluation metrics for simulated data

All peak calling methods were run with the lowest significance threshold possible (p-value or q-value equal to 1 or for any fold enrichment) to generate the complete list of peaks that could be evaluated for their operating characteristics on each of the simulated data sets. Note, GEM only reports exact genomic locations of binding so a 200-bp window around these identified binding locations was used to define peaks for comparison with other methods. For each method, we varied the p-value or q-value threshold to produce a nested set of peaks for evaluating performance across a range of significance levels.

We evaluated performance using several complementary metrics. First, each set of peaks was compared to the location of the binding features in the ChIP sample using the *findOverlappingPeaks* and *annotatePeakInBatch* functions in the ChIPpeakAnno[37] bioconductor[34] package in R[35]. The fraction of the true binding features that overlaps with the significant peaks is defined as the *sensitivity,* the fraction of the significant peaks that overlap with the true binding features is defined as the *precision,* and the harmonic mean of the sensitivity and precision as the *F-score* each method at the particular significance threshold setting on the given simulated data set. We also computed the distance from the center of each significant peak to the center of closest binding feature and used the median of these distances over all significant peaks as a performance metric *(median distance-binding).* We additionally computed the converse metric, the median distance from each of the binding features to the center of the closest significant peak *(median distance-peak).*

The different methods have different typical peak widths, and also the estimated p-values are not always comparable because they result from tests of different hypotheses. Therefore, we chose two approaches to look at the variation of these five metrics *(sensitivity, precision, F-score* and the two *median distance* metrics) as a function of a common measure of a false positive rate across the six methods. First, we used the nested sets of peaks for each method to compare performance at significance levels that produce the same genome coverage (log_10_ fraction of genome). Second, we limited peak lengths to a 200-bp window around either the peak summit or center of the peak (when the information on summit position was not available) for all the methods and compared performance at significance levels that produce the same number of 200-bp peaks. Results were qualitatively the same with both approaches, so we focus on performance as a function of log10 number of peaks.

### Evaluation with Tbx5 ChlP-seq data

We again used the ChIP-seq data sample that measured the binding of the Tbx5 transcription factor to mouse cardiomyocytes(Luna-Zurita et al. Cell 2016). We used the two binding motifs *(Motif 1* and *Motif 2)* [38] given in Figure S2 to represent the *in vitro* sequence binding for Tbx5 in mouse cardiomyocytes. We used the *matchPWM* function in the Bioconductor Biostrings [39] package to identify all the potential binding locations of Tbx5 for each of the two motifs. The threshold likelihood ratio score for defining the potential binding location was fixed at 95% of the maximum possible likelihood ratio value, assuming a zero-order Markov model for a given sequence with prior probabilities for A, C, G and T given by their frequencies in the 2-kilobase (kb) regions of all mouse gene promoters. The shortest genomic distance of each significant peak identified by each peak calling method to binding Motifs 1 and 2 was used as a measure of accuracy. We also compared methods based on the fraction of the top *n* peaks (ordered by statistical significance or by fold enrichment for the thresholding method) that are within 100 bp of Motif 1 and 2.

### Evaluation with H3K36me3 data

ENCODE [40] data for the H3K36me3 histone mark in the GM12878 cell line were used. These files were aligned to the human genome hg19 using Bowtie2[41] and gene counts were obtained using Htseq[42]. We also obtained ENCODE RNA-sequencing data for GM12878 and used edgeR [43] to convert counts to reads per kilobase exon model per million mapped reads (RPKM) and then estimated gene expression by the average of the RPKM values over the two replicates. We considered a peak a positive if it overlaps an active gene (defined varying RPKM from 0 to 2) and compared methods based on sensitivity, precision, and F-score.

### Evaluation with H3K4me3 data

ENCODE[40] data for the H3K4me3 histone mark in the GM12878 cell line were used. We considered a peak a positive if it overlaps the promoter of an expressed gene (RPKM > 0.5). The top 15,000 peak calls from the different methods are ranked by their significance or by their fold enrichment for the thresholding method. We plotted the correct peak fraction (fraction of the top 1000xn peaks that overlap with active promoters) detected as a function of the correct promoter fraction (fraction of the active promoters that overlap with the top 1000xn peaks).

### Binomial test versus the Poisson test

One of the problems that most peak callers need to address is to assign significance to a potential peak region. The significance is based on the rejection of the null hypothesis that the proportion of DNA from a given genomic region in the ChIP sample is less than equal to that in the input sample. This is typically tested by either a Poisson or a Binomial test on the number of reads that map to this genomic location in the ChIP and input sample. We compared the operating characteristics of these two tests using a simulation procedure detailed in the Supplementary Text.

## Results

### Benchmarking peak-calling methods

We benchmarked six peak-calling methods representing different features of the approaches to identifying candidate peaks and evaluating their statistical significance: GEM, MACS2, MUSIC, BCP, TM and ZINBA. These evaluations used 300 simulated and 3 real ChIP-seq data sets. Performance was compared across a range of significance values representing different number of called peaks.

### Simulated Transcription Factor Binding Data

We simulated transcription factor ChIP-seq data with three different noise levels in a manner that closely resembles real data (see Methods). Simulations have the advantage of allowing us to flexibly explore a range of different scenarios in a situation where the ground truth is known.

BCP and MACS2 perform best by sensitivity, precision and F-score metrics across the low (Figure S3), medium (Figure 2) and high (Figure S4) noise levels. TM and ZINBA perform worst, and MUSIC is intermediate. Across methods, except for with ZINBA, median distance of the called peaks to the true peaks and of the true peaks to the called peaks is typically within 100 bp regardless of significance threshold across the low (Figure S5), medium (Figure 3) and high (Figure S6) noise levels. Reduction in simulated noise has the expected effect of improving the sensitivity of BCP, MACS2, GEM and MUSIC, but not of TM and ZINBA (Figure S3). The median distance metrics are predictably larger in the high noise settings (Figure S6).

**Figure 2.**
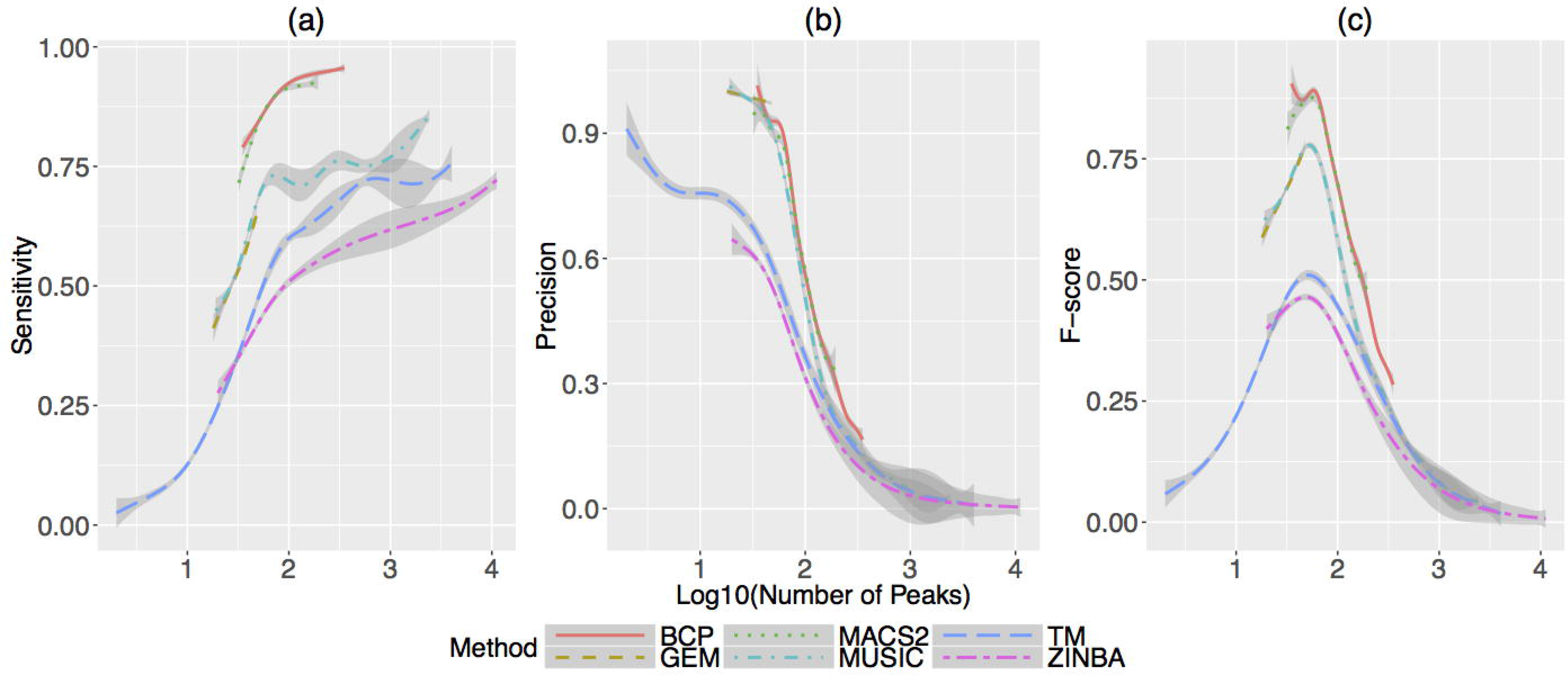
Sensitivity (a), Precision (b) and F-score (c) as a function of the log10 of the number of called peaks for the six peak calling methods on 100 simulated transcription factor ChIP-seq data sets under the medium noise setting. For each method, the means (colored solid lines) and 95% confidence intervals (dark gray regions around the mean profiles) of the means of a metric are estimated using Generalized Additive Models [48] of variation across the 100 simulated data sets, as a function of a smooth function of log_10_ of the number of called peaks.

**Figure 3.**
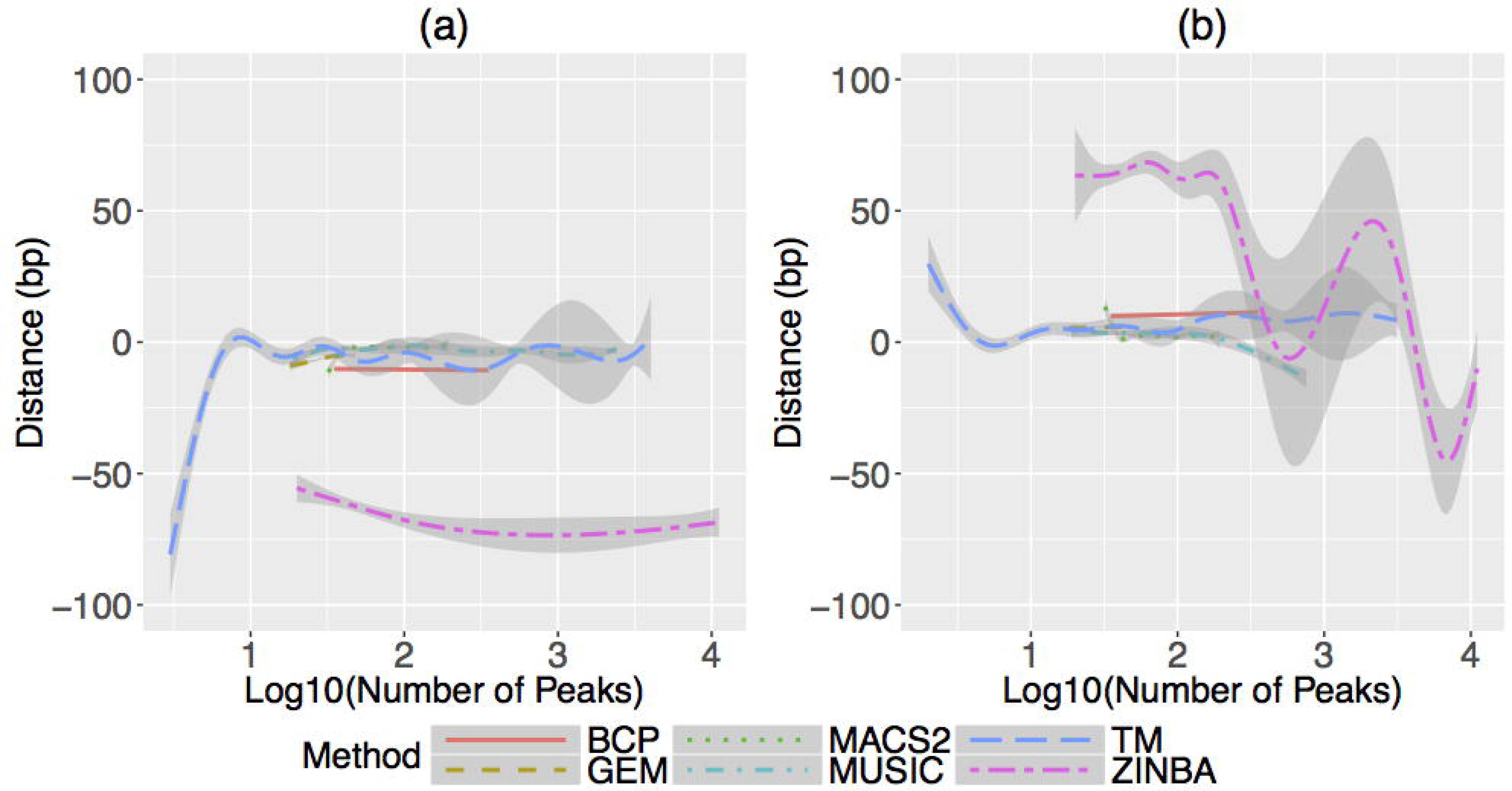
*Median distance-binding* (a) and *Median distance-peak* (b) as a function of the log10 of the number of called peaks for the six peak calling methods on 100 simulated data sets under the medium noise setting. For each method, the means (colored solid lines) and 95% confidence intervals (dark gray regions around the mean profiles) of the means of a metric are estimated using Generalized Additive Models of variation across the 100 simulated data sets, as a function of a smooth function of log_10_ of the number of called peaks.

### Trancription Factor Tbx5 Binding Data

We next evaluated performance of the six methods on data from a Tbx5 ChIP-seq experiment to assess if trends are similar to those revealed by our simulations. Figures 4 and S7 show the fraction of the top *n* peaks that are within 100 bp of a Tbx5 motif. BCP and GEM do particularly well relative to the other methods for Tbx5 Motif 1 (Figure 4). Figures 5 and S8 displays the empirical distribution of the shortest distance of the called peaks to each of the Tbx5 motifs for each method. GEM stands out among all the methods in terms of the fraction of its peaks being closer to a Tbx5 motif than any of the other methods. GEM has the highest fraction of the top 500 peaks with either of two binding motifs of Tbx5, and 50% of its peaks are within 10 bp of a motif (Motif 1; Figure 4). Only 10% of the called peaks of the other methods are within 10 bp of the same Tbx5 motif (Figure 4).

**Figure 4.**
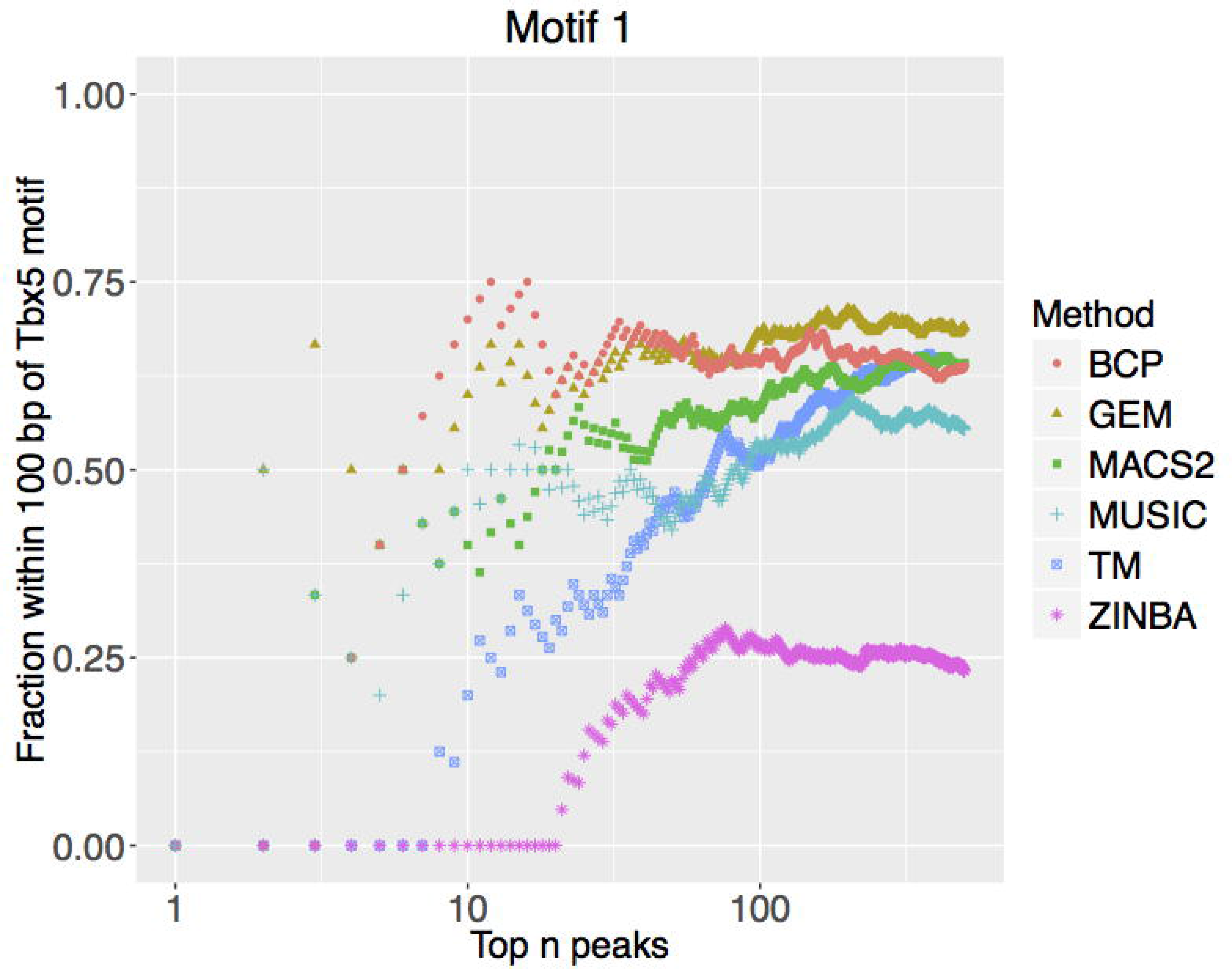
Crypt morphology and differentiation tree. Fraction of top *n* peaks within 100 bp of a Tbx5 motif for the six methods. Results based on Tbx5 Motif 1.

**Figure 5.**
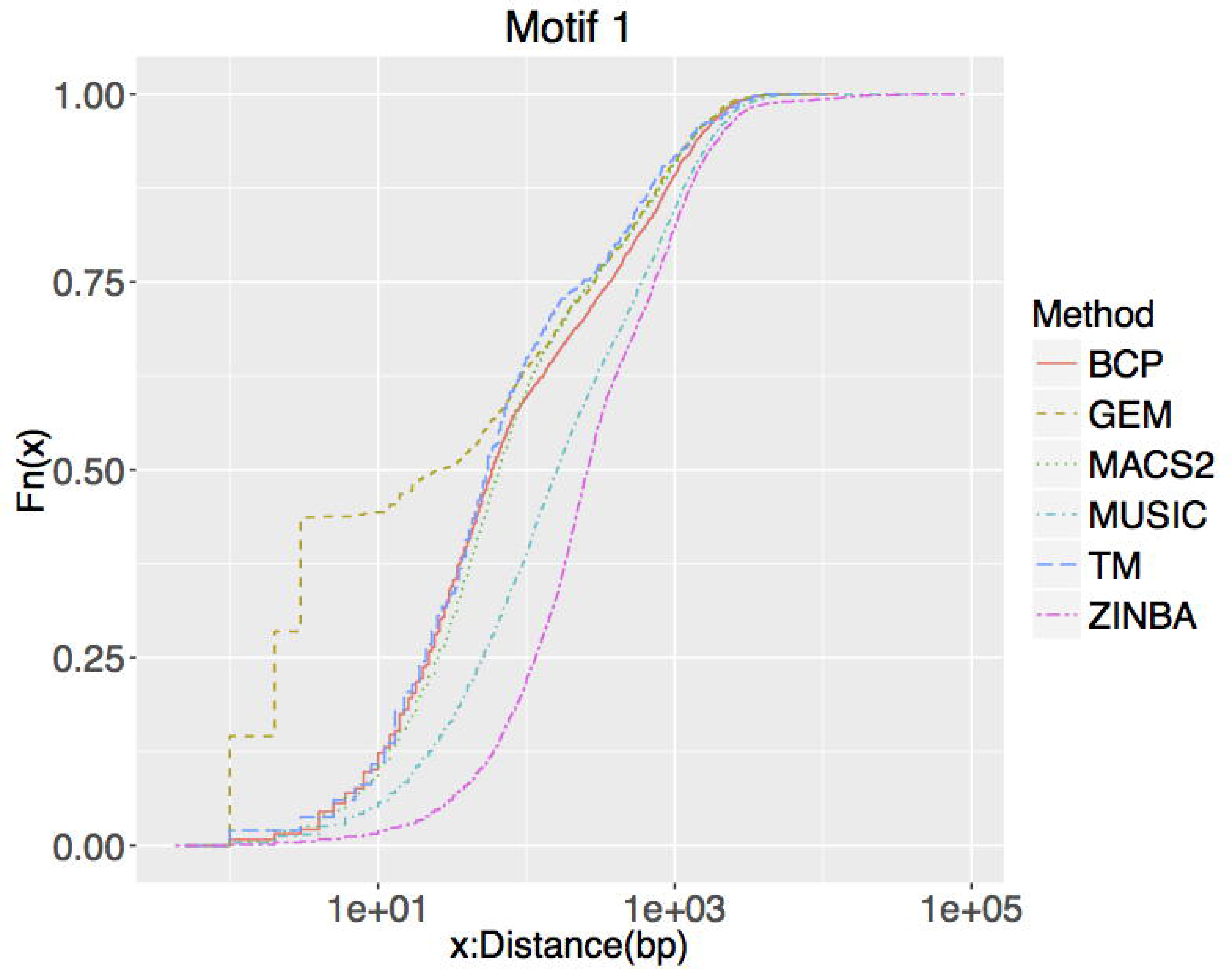
Empirical distribution of the shortest distance to the Tbx5 motif of the significant peaks called by the six methods. Results based on Tbx5 Motif 1.

### Histone H3K36me3 and H3K4me3 Data

Because histones typically have wider peaks than transcription factors and lack DNA motifs, methods perform differently on them. To assess performance on histone ChIP-seq data, we used two sets of experiments from the ENCODE Project [40]. Figure 6 shows the performance of the methods on H3K36me3 data, in terms of how well peaks overlap genes that are actively transcribed (see Methods). MUSIC and BCP perform better than other methods in terms of sensitivity and F-score at a relatively small price in terms of precision. Figure 7 displays the performance of four of the methods on H3K4me3 data, assessed in terms of overlap of peaks with promoters of expressed genes (see Methods). ZINBA failed to run on this data set giving an error that has failed to be resolved with the authors of the software. The other methods perform comparably on this data set, with MUSIC and BCP again being slightly better than the other methods.

**Figure 6.**
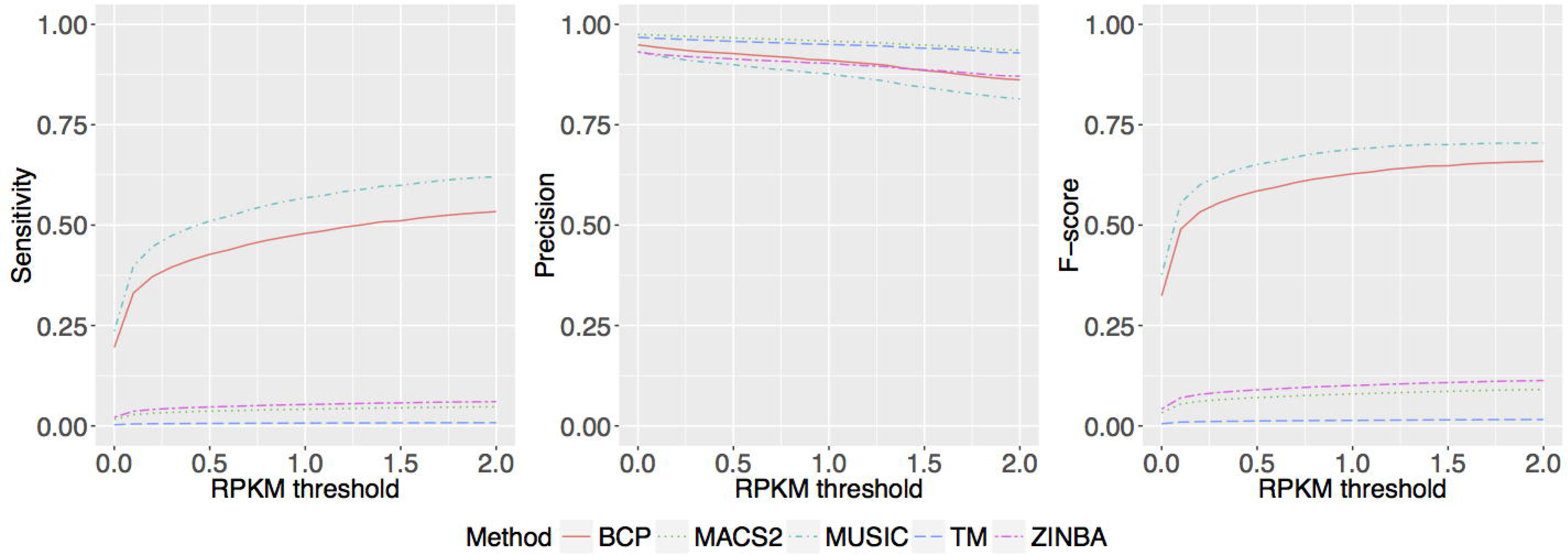
Sensitivity, precision and F-score of the overlap of the called significant peak regions with active gene bodies for H3K36me3 data. The threshold for defining active genes was varied from 0 to 2 RPKM.

### Features of peak calling methods that influence performance

We next investigated which features of peak-calling methods drive the differences in their performance. To do so we identified the features or analysis choices that differentiate the six methods we benchmarked, along with 24 other methods from the literature. Then we evaluated if these features drive performance.

### Sub-Problem 1: Detection of candidate peaks

Four features distinguish methods in terms of how they identify candidate peak regions. Most methods (19 out of 30, Table S1) utilize the signal in a *high-resolution* manner, i.e., peaks could be centered on any nucleotide in the genome as opposed to a bin or a region with more than one nucleotide. There are low-resolution methods (11 out of 30, Table S1) that essentially divide the genome into bins and do not allow signals from one bin to explicitly affect those in the other bins. *ChIP and input sample signal combined* is the second feature that separates methods. Candidate peak regions are defined using the signal that is an explicit combination of the ChIP and input signals either as a fold-change or as a difference score. Methods are deemed not to have this feature (21 out of 30, Table S1) either if they do not use the input signal at all or in cases where the input signal is only used to determine the background rate of reads in the region but is not used to modify the ChIP signal in any way. *Multiple alternate window sizes* (9 out of 30, Table S1) representing widths of influence of each nucleotide for querying the signal could be explicitly or implicitly used. The *variability of the local signal* (18 out of 30, Table S1) is a feature in which the signal is modeled as being generated from a distribution (whose parameters are estimated either using the ChIP or the input signal). Candidate peaks are defined using more than one moment of this distribution and not just the mean.

### Peak detection is reduced when ChlP and input signals are explicitly combined

We mathematically analyzed the probability of detecting a peak when combining ChIP and input data versus not. This analysis was motived by the behavior of TM and ZINBA relative to the other four methods, none of which explicitly combine ChIP and input (Table 1) at the stage of identifying candidate peaks.

Consider a region of the genome with a "true" peak or binding event. Assume that the number of reads from the ChIP sample in this region, denoted by *X* comes from a Poisson distribution with parameter, λ_1_. The number of reads from the input control sample in this region, denoted by *Y* comes from a Poisson distribution with parameter, λ_0_. The reads in the ChIP and input sample are independent of each other and have been normalized for potential differences in sequencing depths. We are interested in identifying this "true" peak. We will compare a measure of the error probability (of the inability to identify this peak) under two scenarios. In the first scenario, only the ChIP signal is used whereas in the second one the ChIP and input signal are explicitly combined as a difference. Let H_0_ and H_1_ denote the null and alternate hypothesis under the two scenarios. (Note: the difference of two Poisson random variables follows a Skellam distribution. The mean and variance of a Poisson random variable with parameter, λ are both equal to λ. The mean and variance of a Skellam distribution with parameters, λ_1_ and λ_0_ are λ_1_-λ_0_ and λ_1_+λ_0_, respectively.)

1. Compare *H*_0_: *X* _˜_ Pois(*λ*_0_) versus *H*_1_: *X* _˜_ Pois(*λ*_1_)
2. Compare *H*_0_: *X - Y* _˜_ *Skellam* (*λ*_1_, *λ*_0_,*λ*_0_) versus *H*_1_: *X - Y* _˜_ Skellam(*λ*_0_,*λ*_1_)

Let *P_e_*(*i, π*) denote the error probability of the inability to distinguish the distributions under H_0_ and H_1_, with prior probabilities *π* = (*π*_0_,*π*_1_) under scenario, *i* = 1,2.

If we can show *P_e_*(1,*π*) < *P_e_* (2,*π*) then using Scenario 1 is preferable to using Scenario 2 to identify the peak. We will use the distance between the underlying distributions of H_0_ and H_1_ as an approximation of this error probability. The intuition is that the farther apart the distributions are the less likely it is to make an error in being unable to distinguish the two distributions. Following Kailath [43], we will use Bhattacharyya distance [44], *B* as a measure of distance between two probability distributions. Let *B*(*i*) denote the distance between the probability distributions associated with H_0_ and H_1_ under scenario, *i* = *1,2*. Kailath [43] showed that if *B*(1) > *B*(2) then there exists prior probabilities, *π* such that *P_e_*(1,*π*) < *P_e_*(2,*π*).

The mean number of reads in the binding regions is typically greater than 20 in a typical experiment [36]. Using this observation and the Central Limit Theorem [45] the normal distribution reasonably approximates both Poisson (Figure S9) and Skellam (Figure S10) distributions. The Bhattacharyya distance between two normal distributions, *N*_1_ and *N*_2_ with parameters, 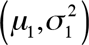 and 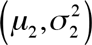 is given by,

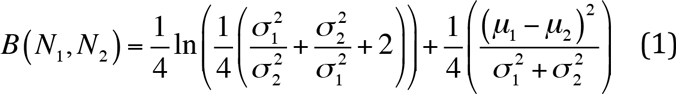

**Theorem 1**: If the Poisson and the Skellam probability distributions can be replaced by Normal distributions with the corresponding means and variances of the respective distributions then there exist prior probabilities, *π* such that *P_e_*(1,*π*) < *P_e_*(2,*π*).

**Proof**: Using equation (1) and the mean and variances for the Poisson and Skellam distributions,

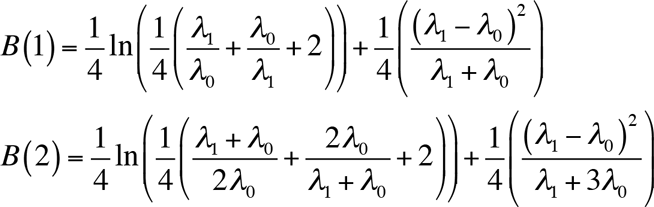

Proving *B*(1) > *B*(2)

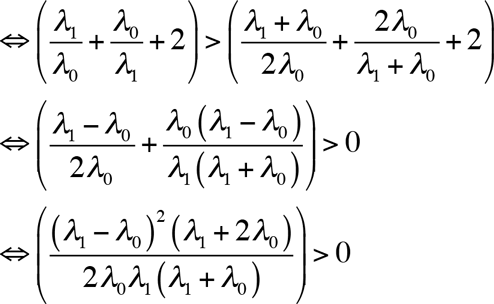

which is true because the by definition the Poisson parameters, *λ*_1_, *λ*_0_ are positive. Using the result in Kailath [43] there exists prior probabilities, *π* such that *P_e_*(1,*π*) < *P_e_*(2,*π*).

Therefore, procedures like TM and ZINBA that explicitly combine the chip and input signals are expected to be less powerful at identifying true binding than ones that do not. This suggests that methods like BCP whose software implementation requires a control sample could in theory be applied to situations where there is no control [46].

### Using input signal to filter candidate peaks

Input is also used to filter regions. GEM implements a filter that removes candidate peak regions with 3-fold or fewer reads in ChIP relative to input, while BCP and MACS2 do not implement a similar filter. We observed that the sensitivity of GEM does not improve beyond a certain level irrespective of how much the significance threshold increases (Figure 2, S3-S4). We hypothesize that filtering may be responsible for this leveling off of performance.

### Window size

Methods use different window sizes to scan the genome for candidate peaks. In our benchmark, TM used a 75-bp sliding window, while MACS2 and MUSIC used 150-bp windows. To check if this difference drives the relative performance of these methods, we implemented MACS2 using window sizes of 75 and 200 bp. The performance of MACS2 with 75-bp windows is worse than with 150-bp and 200-bp windows (Figure S11), suggesting that longer windows are preferable at least for the scenarios we simulated. This raises the question of whether the optimal window size is different for narrow versus broad peaks. To explore this, we mathematically analyzed variation in the likelihood of detecting a peak of length *l* using a window size *w* to scan the genome.

Let *γ*_1_ be the Poisson rate parameter for the number of reads per base in the peak corresponding to a binding event. So the number of reads in a w-bp wide interval in the peak is distributed as a Poisson random variable with parameter, *γ*_1_^w^. Also, let *γ_0_* be the Poisson rate parameter for the number of reads per base in the input sample. Let *X* represent the number of reads arising from a window of width *w* bp.

Consider two situations, *w ≤ l* and *w > l*,

For the situation *w ≤ l*

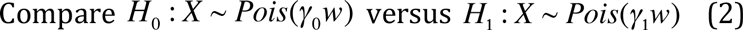

For the situation *w > l*

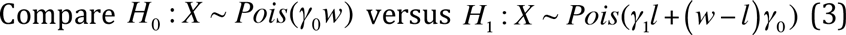

We will again use the Bhattacharyya distance as a measure of the ability to distinguish between the two distributions in Equations (2) and (3). The Bhattacharyya distance between two Poisson distributions of rate parameters, (*λ*_1_, *λ*_2_) is given by,

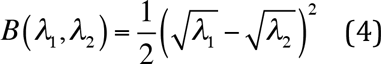

Let *B*(*w*) denote this distance measure as a function of the size *w* of the window used to scan the genome. Then using the Poisson rate parameters in Equations (2) and (3) and the formula for distance in Equation (4),

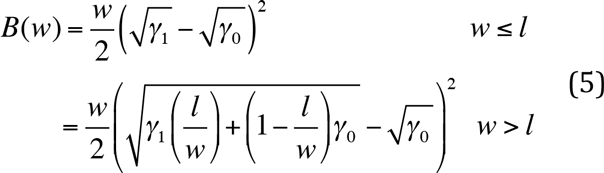

Therefore, the distance between the probability distributions associated with the null and alternate hypothesis increases linearly with *w* until it reaches the actual peak length and thereafter decreases asymptotically to zero. Denote the error probability of the inability to distinguish the two signals in the peak and background as *P_e_*(*w, π*) that is a function of *π* and prior probabilities *π* for the two hypotheses. Using the result from [43], there exist prior probabilities *π* such that the value of *P_e_*(*w, π*) decreases to a minimum when *w* = *l* after starting from a window of size one base and then again increases asymptotically to 1 with increasing window width (Figure S12).

This result that the optimal window size for scanning the genome is the true peak width suggests an explanation for the performance results we observed with the histone data (Figures 6-7). Harmanci et al [7] present an estimate of the spectrum of peak lengths associated with a transcription factor and different histone marks including H3K36me3 and H3K4me3. A characteristic of most of these spectra is the presence of peaks that vary in lengths across orders of magnitudes. Therefore methods that work only with one window size are biased to pick peaks of length only of comparable magnitude.

**Figure 7.**
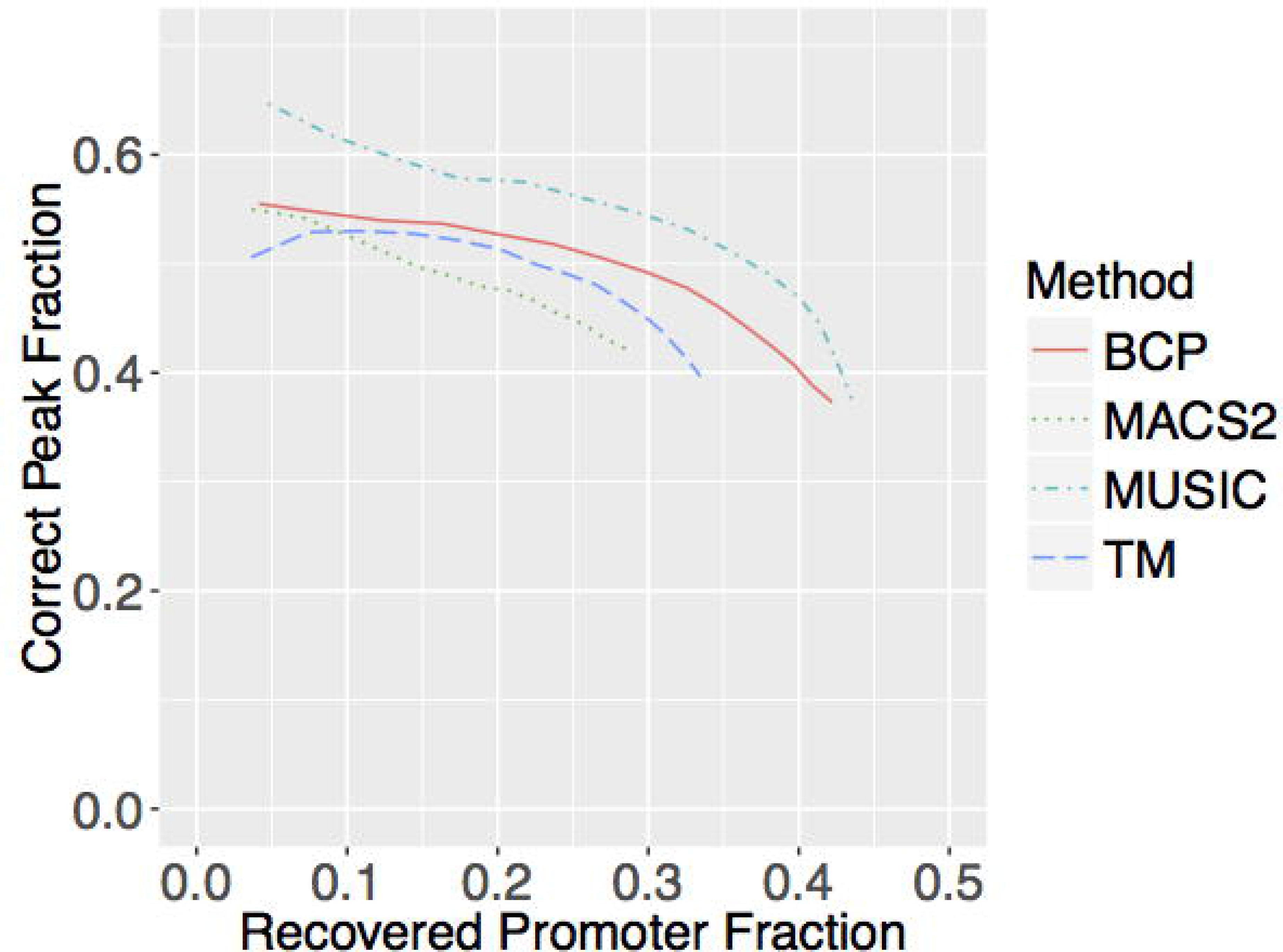
Correct peak fraction (fraction of top called 1000n peaks that overlap with the promoters of active genes, genes with expression > 0.5 rpkm) as a function of recovered promoter fraction (fraction of promoters of active genes that overlap with the top called 1000n peaks) for H3K4me3 data. The peaks for each method were ranked by their assigned significance.

### Incorporating variability of the local signal

The difference in the operating characteristics of MUSIC as compared with MACS2 and BCP can potentially be explained by the fact that in identifying their candidate peak regions MACS2 and BCP uses the variability of the local signal while MUSIC does not. The peak regions are identified by local minima of the smoothed signals by MUSIC, whereas MACS2 checks if the number of reads in a candidate region is different from expected assuming a given background rate using a Poisson test, and BCP implicitly takes into account the variability of the local signal by its estimation of the parameters associated with its hidden Markov model.

### Sub-Problem 2: Statistical significance of candidate peaks

Once the candidate peaks have been identified, the different methods typically rank candidates by their significance of a hypothesis test that compares the counts in the corresponding genomic regions of the ChIP and input samples. This test has mostly been implemented as a *Poisson* (9 out of 30, Table S1) test, a *Binomial* (7 out of 30, Table S1) test or by *fold-change or Normalized Difference* (5 out of 30, Table S1). There are methods that rank the peaks by some *posterior* measure (6 out of 30, Table S1) that could be the posterior probability of binding at a given genomic region or the posterior rate of counts in a given genomic region. Additionally, there are methods that explicitly use the underlying genome sequence (1 out of 30, Table S1) or the shape of the candidate peaks (2 out of 30, Table S1) in assigning significance values.

### Binomial versus Poisson Test

BCP and MACS2 have the best operating characteristics on simulated data (Figures 2, S3-S4). Could this be because of how these methods test candidate peaks for statistical significance? BCP and MACS2 use the Poisson test, while MUSIC and GEM use the Binomial test. To explore this question, we first used simulations (see Methods) to directly compare Poisson and Binomial tests on a predefined set of candidate peaks. These empirical results show that the Poisson test is more powerful at detecting enriched regions, while maintaining a reasonable Type I error rate (Figure 8). Second, we implemented modified versions of BCP and MUSIC that expand the methods to allow either a Binomial or Poisson test. Note our Poisson test version of BCP and Binomial test version of MUSIC are essentially the same as in the original methods. In this approach, we are using the read counts in the regions called peaks by each method and performing the statistical test in two ways. The operating characteristics of both versions of each method are hardly distinguishable (Figures S13-S14). However, in real situations one works with a set of peaks identified by a chosen threshold. Analyzed this way, rather than over all possible thresholds, the Poisson version of each method is clearly better (Figures S15-S16).

**Figure 8.**
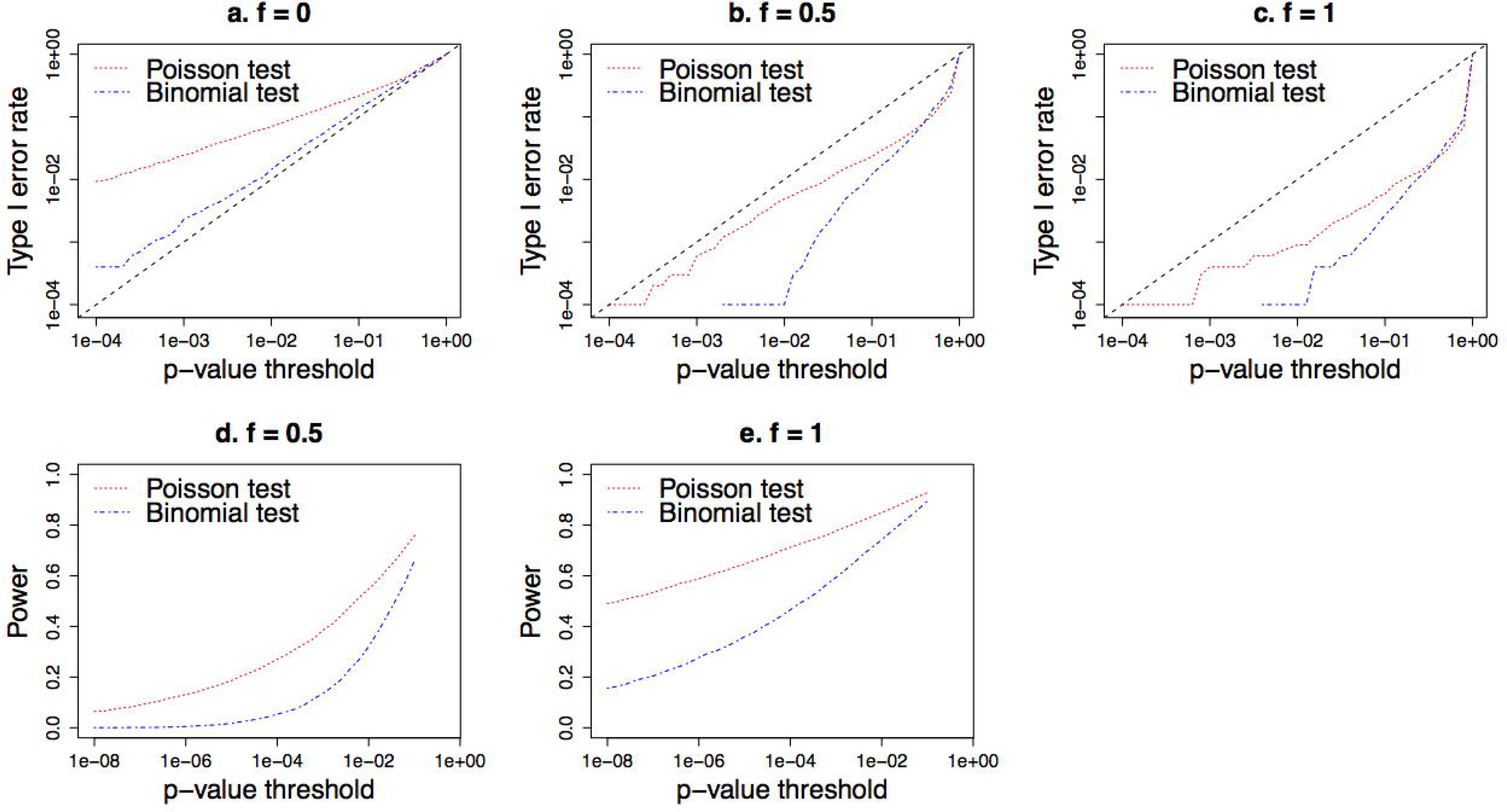
Type I error rate and statistical power comparison between Poisson and Binomial tests. *f* is a parameter that controls the increase in the proportion of DNA from a given region in the input relative to the ChIP sample for the Type I error evaluations and increase in this proportion for the ChIP relative to the input sample for the power evaluations.

## Discussion

We performed a benchmarking study and systematic evaluation of the features of ChIP-seq peak calling methods that drive their relative performance.

Our benchmarking analysis included six commonly employed methods that are representative of the different features of ChIP-seq software tools. Overall, BCP and MACS2 have the best operating characteristics on simulated transcription factor binding data. On real Tbx5 ChIP-seq data, GEM stood out in terms of how close its peaks are to the primary binding motif of Tbx5. BCP and MUSIC perform best on histone ChIP-seq data.

We identified three features that are the most important drivers of performance in peak calling. First, methods that explicitly combine ChIP and input signal in trying to identify candidate peak regions are less powerful than those that only use the signal from the ChIP sample. Second, methods that use windows of multiple widths to scan the genome for candidate peaks perform better than others. This is particularly important with broad histone marks. Use of multiple window sizes can be implemented either explicitly (e.g., MUSIC) or implicitly (e.g., BCP via a hidden Markov model), but most methods we examined used only a single size. Finally, the Poisson test is more powerful than the Binomial test for statistically scoring candidate peaks.

A few other features of peak-calling methods merit consideration. Our results suggest that methods utilizing variability of the local signal in identifying candidate peak regions are likely to have better operating characteristics than ones that do not. Secondly, the ability of GEM to call peaks very close to motifs of the immunoprecipitated transcription factor points to the benefit of incorporating the underlying genome sequence and knowledge of binding sites at the stage of ranking candidate peaks. Note that there are other methods of ranking candidate peak regions based on their shape characteristics [22,24] that haven’t been evaluated in this manuscript, which may also provide performance benefits.

This manuscript focuses is on how peak calling methods differ in terms of how they identify candidate peaks and compute their statistical significance. All peak calling methods sequentially implement solutions to these two problems, and then most use a significance threshold to determine a set of peak calls [47]. We note that selecting a threshold is reasonably straight forward; most methods use a false-discovery rate based multiple testing-correction and a user defined proportion of false discoveries that will be tolerated in a given application. We therefore did not compare methods based on a single choice of significance threshold, but instead examined performance across a range of thresholds.

## Acknowledgements

This work was supported by the NHLBI Bench-to-Bassinet program (grant #HL098179), NHLBI grant #HL089707, and BioFulcrum: A Gladstone Institutes Enterprise.

